# The number of lateral hypothalamus orexin/hypocretin neurons contributes to individual differences in cocaine demand

**DOI:** 10.1101/547836

**Authors:** Caroline B. Pantazis, Morgan H. James, Brandon S. Bentzley, Gary Aston-Jones

## Introduction

Orexin A and B (also called hypocretin 1 and 2) are peptides synthesized by a small population of neurons in posterior hypothalamus, ranging mediolaterally from dorsomedial hypothalamic nucleus (DMH), through perifornical hypothalamus (Pef), to lateral hypothalamus (LH) (de Lecea, Kilduff, Peyron et al., 1998; Sakurai, Amemiya, Ishii et al., 1998). Orexin A binds both orexin-1 and −2 receptors (OxR1 and OxR2), whereas orexin B preferentially binds OxR2 (Zhu, Miwa, Yamanaka et al., 2003). Orexin neurons interact closely with the mesolimbic dopamine (DA) system to drive motivation and addiction (Borgland, Taha, Sarti et al., 2006; Espana, Oleson, Locke et al., 2010; Fadel & Deutch, 2002; James, Charnley, Levi et al., 2011; Mahler, Moorman, Smith et al., 2014; Mahler, Smith & Aston-Jones, 2013; Mahler, Smith, Moorman et al., 2012). Many studies now indicate that blockade of OxR1 signaling attenuates cocaine seeking behaviors, particularly under high effort conditions (for review, (James, Mahler, Moorman et al., 2017), indicating a role for OxR1 signaling in the motivational properties of cocaine.

A major goal of addiction research is to elucidate the neural substrates that contribute to addiction vulnerability. We previously found that the degree to which drug-associated cues and contexts activate the LH subpopulation of orexin neurons predicts individual differences in drug seeking. Fos activation of LH, but not Pef/DMH, orexin neurons correlated with reinstatement of extinguished cocaine or morphine preference (Harris, Wimmer & Aston-Jones, 2005). LH orexin Fos expression also correlated with preference for, and context-induced reinstatement of, ethanol seeking (Moorman, James, Kilroy et al., 2016). Further, rats that exhibited a multifaceted addiction phenotype following intermittent access (IntA) to cocaine exhibited higher Fos expression selectively in LH orexin neurons following re-exposure to the self-administration context relative to short access control rats (James, Stopper, Zimmer et al., 2018). OxR1 blockade is particularly effective at reducing drug-seeking in highly motivated animals (James, Bowrey, Stopper et al., 2018; James, Stopper, Zimmer et al., 2018; Jupp, Krstew, Dezsi et al., 2011; Lawrence, Cowen, Yang et al., 2006; Lopez, Moorman, Aston-Jones et al., 2016; Moorman & Aston-Jones, 2009; Moorman, James, Kilroy et al., 2017), further indicating that individuals with high propensity for drug seeking may have elevated stimulus-driven LH orexin cell activity.

Although it is clear that the activity of LH orexin neurons is important for drug seeking, emerging evidence indicates that upregulation of orexin expression may also be involved. We reported that the persistent IntA-induced addiction phenotype was associated with higher numbers of orexin-expressing neurons in LH but not Pef/DMH (James, Stopper, Zimmer et al., 2018). Another recent study found that postmortem tissue from heroin addicts contains higher numbers of orexin-expressing neurons compared to healthy controls, and mice exposed to chronic non-contingent morphine injections had increases in the number of orexin neurons particularly in LH (Thannickal, John, Shan et al., 2018). These studies are consistent with an earlier study that reported that chronic alcohol consumption increased the area of prepro-orexin mRNA expression preferentially in LH of alcohol-preferring rats (Lawrence, Cowen, Yang et al., 2006). Changes in the number of orexin-expressing LH neurons might reflect individual differences in drug seeking that result from ‘state’ factors like extended or binge-like drug access. However, it is unclear whether variability in the baseline (non-stimulated) numbers of orexin-expressing LH neurons contributes to trait differences in addiction behavior.

We sought to determine if differences in the numbers of LH orexin neurons contributes to individual differences in baseline or ‘trait’ motivation for cocaine using our within-session behavioral economics (BE) paradigm (Bentzley, Fender & Aston-Jones, 2013; Bentzley, Jhou & Aston-Jones, 2014). In this paradigm, cocaine consumption is measured at increasing price points. By fitting the resulting data using an exponential demand equation (Hursh & Silberberg, 2008), it is possible to calculate demand elasticity (α), or the extent to which consumption changes with price. We previously reported that α predicts several addiction-relevant behaviors, including compulsive (punished) responding for cocaine, drug seeking during initial abstinence, and cue-induced reinstatement of extinguished drug seeking (Bentzley, Jhou & Aston-Jones, 2014; James, Bowrey, Stopper et al., 2018). Here, we show that the number of orexin neurons in LH, but not Pef/DMH, correlates with motivation for cocaine (α), and this relationship persists after abstinence. In addition, unilateral LH orexin knockdown with orexin-A morpholino antisense is sufficient to increase demand elasticity (reduce motivation), and the degree of knockdown predicts the extent of motivation reduction. Collectively, these studies indicate that the number of LH orexin neurons is a potent predictor of addiction vulnerability.

## Methods

### Animals

Adult male Sprague-Dawley rats (300-325 grams) were pair-housed on a reverse 12-hour light:dark cycle in a temperature- and humidity-controlled animal facility at Rutgers University or Medical University of South Carolina (MUSC) with *ad libitum* access to standard rat chow and water. Upon arrival, animals were acclimated to the colony room for 2 days and handled for at least 3 days prior to surgery. All protocols and animal care procedures were approved by the Institutional Animal Care and Use Committee at Rutgers University-New Brunswick or Medical University of South Carolina

### Drugs

Cocaine HCl powder was obtained from the National Institute of Drug Abuse (Research Triangle Park, NC) and dissolved in 0.9% sterile saline.

### Intravenous catheter surgery

After handling for a minimum of 3 days, rats were anesthetized with a ketamine/xylazine (56.5/8.7 mg/kg, i.p., respectively) mixture and also given an analgesic (rimadyl at 5 mg/kg or meloxicam at 1 mg/kg, s.c.). Rats were implanted with indwelling catheters into the jugular vein for iv infusion of cocaine. For morpholino experiments, cannulae implantation occurred immediately followed catheterization. Cefazolin (0.1 ml; 100 mg/ml) and heparin (0.1 ml; 100 U/ml) were flushed through the iv catheter after surgery, and after each self-administration session. Following a one-week recovery after surgery, animals were trained to self-administer iv cocaine.

### Cocaine self-administration

Rats were trained on a fixed ratio-1 (FR-1) cocaine self-administration paradigm (20-second timeout post-infusion) as previously described (McGlinchey, James, Mahler et al., 2016). Sessions occurred in operant chambers contained in sound-attenuating boxes with Med-PC IV software (Med Associates). During training sessions, cocaine infusions (0.19 mg cocaine/infusion) were paired with discrete light and tone cues (white stimulus light above the active lever; 78-dB, 2900-Hz tone). After reaching criteria, (≥ 10 infusions/session for 10 sessions), animals were trained on a BE procedure.

### Behavioral Economics

Following FR-1 training, rats were trained on a within-session BE procedure, as described previously (Bentzley, Fender & Aston-Jones, 2013; Bentzley, Jhou & Aston-Jones, 2014). During the 110-minute session, the price of cocaine increased in successive 10-minute intervals on a quarter logarithmic scale (383.5, 215.6, 121.3, 68.2, 38.3, 21.6, 12.1, 6.8, 3.8, 2.2, 1.2 μg cocaine per infusion). Lever pressing responses were fit to a demand curve, as described in a previous paper from our lab (Bentzley, Fender & Aston-Jones, 2013). From the demand curve, we derive consumption at low effort (Q_0_) and demand elasticity (α, the rate of decline in consumption as price increases). Q_0_ is low effort consumption extrapolated to the y-axis of the demand curve, where cocaine price approaches null; α is the decay constant of this curve. Thus, α scales inversely with motivation, so high motivation animals have lower α values. Animals were trained on BE for a minimum of six days. Testing occurred when animals displayed stable behavior on the BE paradigm (Q_o_ and α values ≤ 30% variability across the last three sessions).

### Tissue preparation for immunohistochemistry

Animals were deeply anesthetized with a ketamine/xylazine mixture and transcardially perfused with 0.9% sterile saline then 4% paraformaldehyde. Brains were dissected and postfixed in 4% paraformaldehyde, then 20% sucrose-PBS azide solution. Brains were frozen with dry ice and sectioned at 40 um on a cryostat. Sections were collected in PBS azide.

### Immunohistochemistry

To visualize orexin neurons, hypothalamic tissue was incubated in goat anti-orexin A (1:500, Santa Cruz Biotechnology) in 5% NDS at room temperature overnight. The following day, tissue was incubated in biotinylated donkey anti-goat secondary antibody (1:500, Jackson ImmunoResearch Laboratories) for 1.5 hours, followed by avidin-biotic complex (1:500) for 1.5 hours. After washing, sections were stained with DAB (Sigma) in Tris buffer. Tissue was mounted on slides, dehydrated, and coverslipped with DPX mounting medium (Electron Microscopy Sciences).

### Cell quantification

Coronal lateral hypothalamus images were taken across the orexin cell field (2.5-3.8 mm caudal to bregma) with a Zeiss Axio Zoom V16 microscope (Paxinos and Watson, 2007). Tiled photographs were compiled at 20x magnification using Zen 2 imaging software (Carl Zeiss Microscopy). Orexin A- and Fos-immunopositive neurons were quantified in both hemispheres with Adobe Illustrator by an observer blind to experimental conditions. Three sections were taken per animal, as in our previous studies (James, Stopper, Zimmer et al., 2018; Mahler & Aston-Jones, 2012; Moorman, James, Kilroy et al., 2016). The sum of neurons counted in both hemispheres in each section were determined and averaged across the three sections. To determine the extent of orexin A knockdown, the percent change in injected vs. non-injected hemispheres for each section was averaged across the three sections.

#### Experiment 1: Relationship between orexin cell numbers and economic demand for cocaine

We first sought to examine the relationship between endogenous orexin cell numbers and baseline economic demand for cocaine. To do this, animals were trained on the BE paradigm as above until they displayed stable behavior. The following day, rats were again tested on the BE paradigm and perfused 90 min after the point of maximum responding (Pmax), defined as the price at which maximum responding occurred. Tissue was processed for orexin A immunohistochemistry and cell quantification; cell counts were compared to demand measures from the final BE test session.

#### Experiment 2: Effect of orexin cell knockdown on economic demand for cocaine

##### Stereotaxic surgery

We next investigated the impact of knocking down LH orexin expression using orexin-A morpholino antisense on demand for cocaine. Immediately following catheter implantation (described above), animals were secured in a stereotactic frame (Kopf, Tujunga, CA, USA) and implanted with a unilateral stainless steel guide cannula (22 gauge, 11 mm, Plastics One, Roanoke, VA, USA) 2 mm dorsal to LH (coordinates relative to bregma: −3.0 mm AP, +/-2.6 mm ML, −6.8 to −7.4 mm DV). We varied the dorsal-ventral cannula coordinates to produce varying degrees of knockdown in morpholino-infused animals. The implanted hemisphere was counterbalanced across all tested animals, such that an equal number of animals were implanted in the left vs. right hemisphere. Acrylic cement and jeweler screws were used to secure the cannula to the skull surface.

##### Orexin morpholino antisense microinjections

One day after animals displayed stable behavior on the BE paradigm, an injector cannula was unilaterally lowered into LH (2 mm below the guide cannula) to infuse orexin-A morpholino antisense (Vivo-Morpholino; 150nmol/0.3 uL in 0.5mM phosphate buffer, Gene Tools, 5’-GTATCTTCGGTGCAGTGGTCCAAAT-3’) or an inverted orexin control missense (Vivo-Morpholino, 5’-TAAACCTGGTGACGTGGCTTCTATG-3’). The injector cannula was kept in place for 1 minute before infusing the morpholino for 2 min. Injectors were then kept in place for 1 minute to reduce diffusion of the morpholino. All animals were infused between 14:00-15:00h, during their active period. Animals were tested on BE for 6 d following morpholino microinjection, and sacrificed perfused on day 6 when peak orexin knockdown occurs (Reissner, Sartor, Vazey et al., 2012; Sartor & Aston-Jones, 2012). Animals were perfused with fixative, brains were sectioned, sections were stained for the orexin A protein, and orexin-positive cells were quantified as above. By restricting morpholino injections to a single hemisphere, we were able to simultaneously: i) confirm the efficacy of the antisense by comparing the number of orexin cells in the injected and uninjected hemispheres; ii) examine the effect of reducing overall orexin numbers on demand for cocaine; and iii) examine the relationship between endogenous orexin cell numbers (in the uninjected hemisphere) with demand prior to antisense injections.

Brain sections adjacent to those used for orexin A staining were dehydrated and Nissl-stained with neutral red. Sections were coverslipped with DPX mounting medium (Electron Microscopy Sciences) to determine cannula location.

#### Experiment 3: Relationship between LH orexin cell number and demand following abstinence

To determine whether the relationship between LH orexin cell numbers and demand persisted after abstinence, we sacrificed animals with a history of cocaine self-administration two weeks after their final BE test. Animals were first trained on the BE procedure until behavior was stable, as above. Cocaine self-administration was then discontinued, and rats were then tested for locomotor activity in locomotor chambers (clear acrylic, 40 x 40 x 30 cm) equipped with Digiscan monitors (AccuScan Instruments) in 2 h sessions, 5d/wk for 2 wk. During these 2 wks, rats received ip injections of up to 3 compounds (propranolol, prazosin, clonidine) as a control for a separate study. At least 2d washout was given between treatments and before the final sacrifice. Horizontal, vertical, and total locomotor activity was recorded using beam beaks. Rats were perfused immediately after the final habituated locomotor test for orexin cell quantification.

##### Data analysis

Statistical analyses were performed with GraphPad Prism 7, except for multivariate regressions, which were performed using SPSS Statistics (Version 22). Baseline α and Q_0_ values were determined by averaging performance across the three days preceding testing. In Experiment 1, a median split of α was used to determine high demand elasticity vs. low demand elasticity animals, and a median split of Q_0_ identified high takers vs. low takers. Unpaired samples *t*-tests were used to compare differences in LH orexin neuron numbers between high and low demand elasticity, or high and low Q_0_ animals. In Experiment 2, a two-way ANOVA was used to compare BE performance following morpholino treatments. Demand associations with orexin cell counts were determined using multiple linear regression with log10(α) and log10(Q_0_) set as independent variables with cell count as the dependent variable. In Experiment 3, an unpaired samples *t*-test was used to compare LH orexin neuron numbers in animals subjected to BE vs. locomotor testing. Pearson correlations were used for all univariate relations to compare orexin neuron numbers with total locomotor activity or α values. All statistics were two-tailed. A Shapiro-Wilk normality test determined that all data were parametric.

## Results

### Experiment 1: Animals with high motivation for cocaine have more LH orexin-expressing neurons

Animals (n=12) were trained on the BE paradigm and divided into high vs. low demand elasticity based on a median split of α values (representative demand curves shown in Figure 1A). Low demand elasticity (high motivation) animals had significantly lower α values than high demand elasticity (low motivation) animals (Figure 1B, t_10_=3.53, *p*<0.01). Low demand elasticity animals also had more orexin-immunopositive cells in lateral hypothalamus (LH), compared to high demand elasticity animals (Figure 1D, t_10_= 2.32, *p*<0.05). No such differences were observed for perifornical area/dorsomedial hypothalamus (Pef/DMH) orexin neurons (Figure 1E, t_10_=1.138 *p*=0.282). When animals were instead divided based on their preferred level of consumption at low effort (Q_0_), there was no difference in the number of LH (Figure 1F, t_10_=0.29, *p*=0.776) or Pef/DMH (Figure 1G, t_10_=0.068 *p*=0.947) orexin neurons between high- and low-takers.

**Figure 1.**
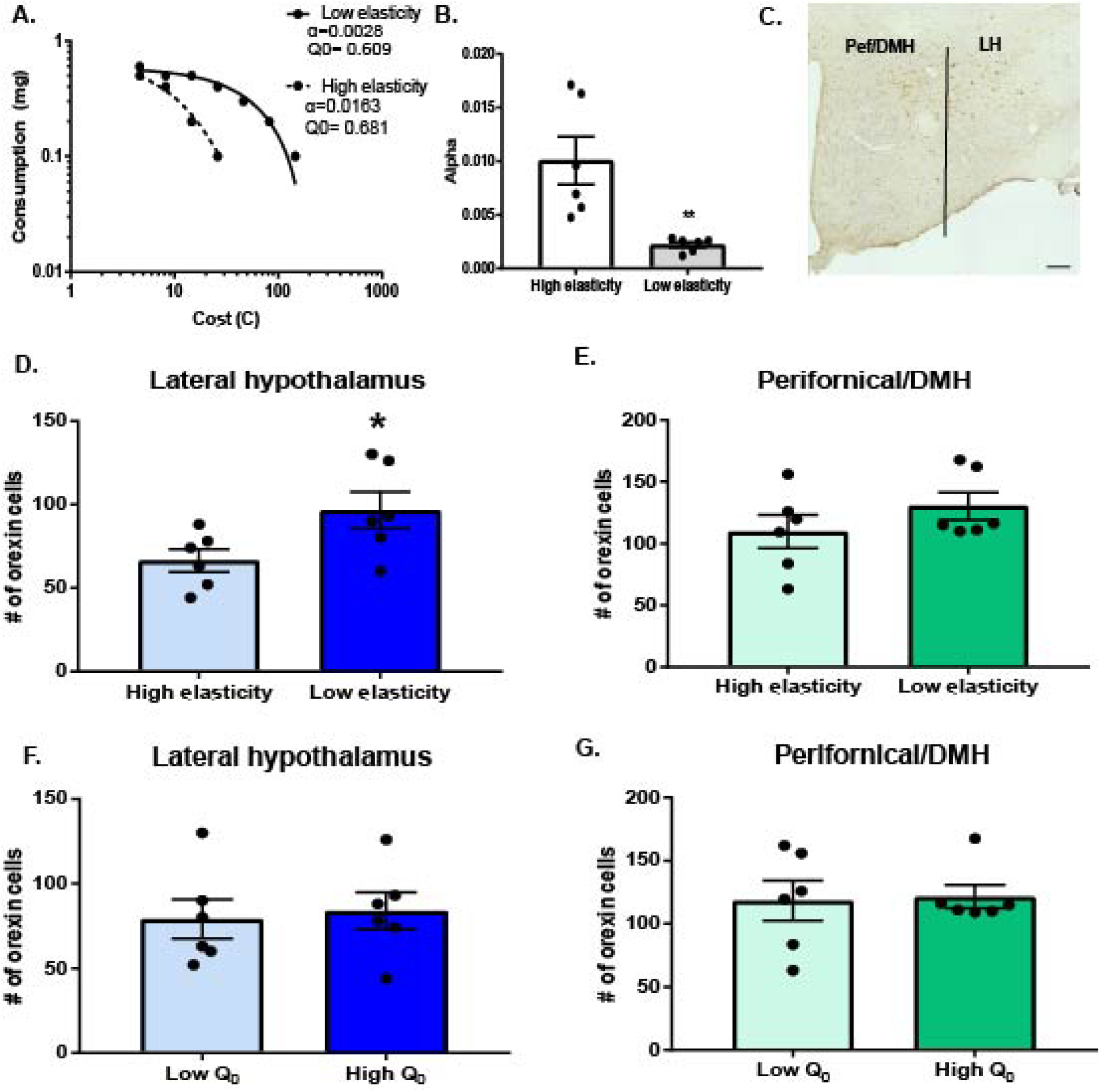
Number of LH orexin neurons is elevated in low demand elasticity (high motivation) animals. <u>A.</u> Representative demand curves from high (dashed line) and low (solid line) demand elasticity animals with similar baseline consumption (Q_0_). <u>B</u>. Low demand elasticity (high motivation) animals had significantly lower α values than high demand elasticity animals (unpaired samples t-test, \*\**p*<0.01). <u>C</u>. Representative section of hypothalamus stained for orexin from an animal sacrificed 90 minutes post-Pmax. Scale bar denotes 200 µm. <u>D</u>. Low demand elasticity (high motivation) animals had more LH orexin neurons, compared to high demand elasticity (low motivation) animals (n=6/group, unpaired samples t-test, \**p*<0.05). Bilateral orexin neurons in Pef/DMH and LH were counted separately and averaged across three sections per animal, representing the rostral-caudal extent of the orexin cell field for this panel and for panels E-G. <u>E</u>. High and low demand elasticity animals had similar numbers of Pef/DMH orexin cells (unpaired samples t-test, *ns*). <u>F</u>. No differences were observed in LH orexin numbers in high Q_0_ compared to low low Q_0_ (unpaired samples t-test, *ns*). <u>G</u>. High and low Q_0_ animals did not differ in the number of Pef/DMH orexin cells (unpaired samples t-test, *ns*).

### Experiment 2: Unilateral knockdown of LH orexin neurons reduces cocaine demand elasticity

We next examined whether the number of LH orexin neurons plays a causal role in demand by testing whether LH orexin knockdown would reduce motivation for cocaine (increase α). In a separate cohort of animals, we unilaterally infused an orexin-A morpholino antisense (n=9) or control missense (n=8) into LH after animals displayed stable BE behavior. Injection sites are depicted in Figure 2A.

**Figure 2.**
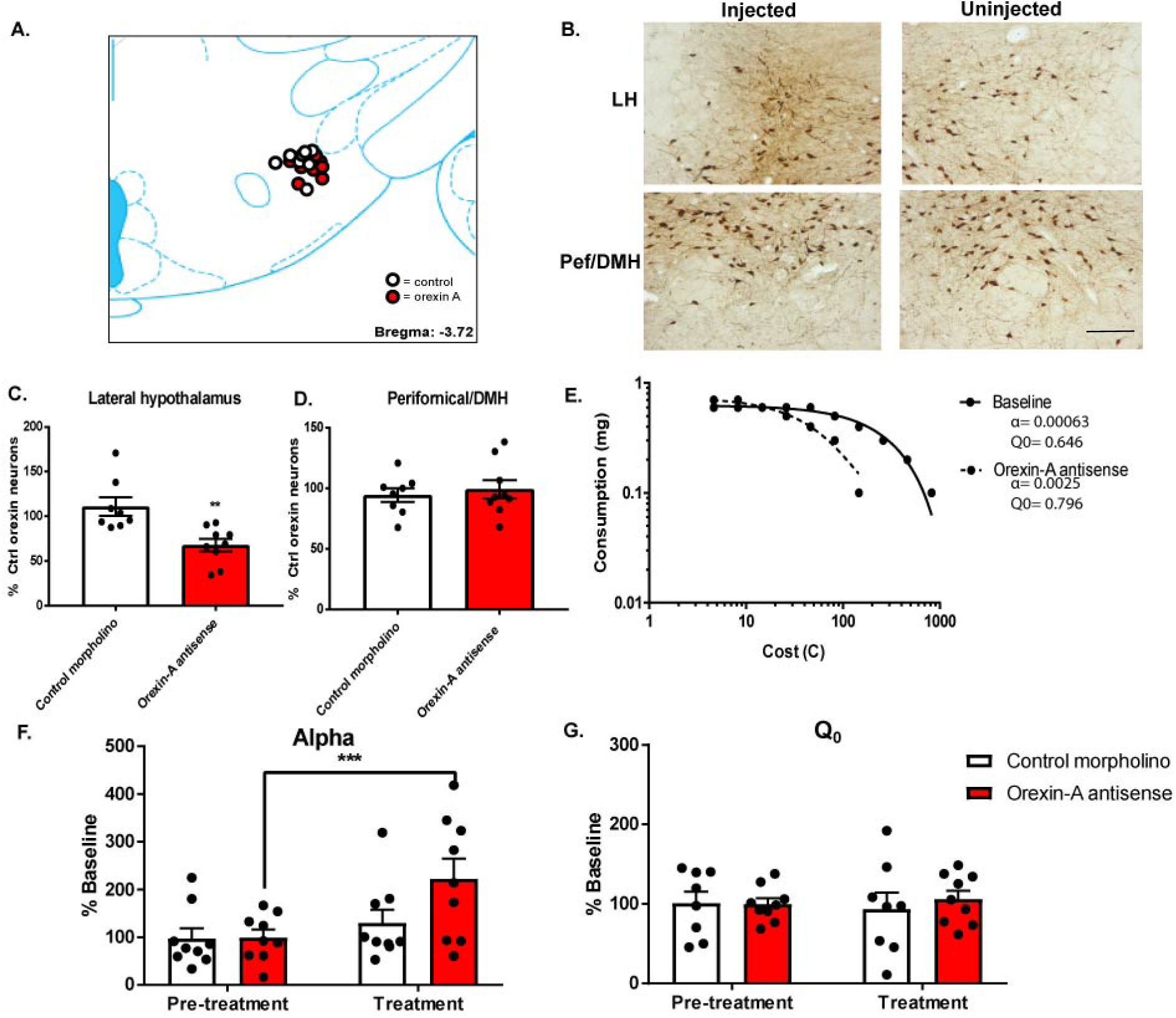
Knockdown of LH orexin-expressing neurons with unilateral morpholino antisense decreases demand for cocaine. <u>A.</u> Schematic representation of LH cannula placements for control missense-(n=8, white circles) and orexin-A antisense-(n=9, red circles) infused animals. <u>B.</u> Representative images of hypothalamus stained for orexin-A (brown) following a unilateral orexin-A antisense-infusion. The uninjected hemisphere was used as a within-subjects control. The top left panel indicates LH orexin cells in the injected hemisphere, top right LH orexin cells in the uninjected hemisphere. The bottom left panel indicates Pef/DMH cells in the injected hemisphere, whereas the bottom right panel shows those cells in the uninjected hemisphere. Scale bar denotes 200 µm. <u>C.</u> The percent reduction in orexin expression in orexin-A antisense-infused animals was significantly greater than in the control missense-infused animals (unpaired t-test, *p*<0.01). <u>D.</u> Pef/DMH orexin expression was unaffected in orexin-A antisense and control missense-infused animals, compared to the uninjected hemisphere (unpaired t-test, *ns*). <u>E.</u> Sample demand curves for an antisense-infused animal at baseline (solid line) and following morpholino treatment (dashed line). <u>F.</u> Microinjection of orexin-A antisense increased α (decreased motivation) compared to baseline (red bars; Sidak’s comparisons test, \*\*\**p*<0.001). No change was observed in control missense-infused animals (white bars; *ns*). <u>G.</u> There was no effect of treatment on Q_0_ in orexin-A antisense-(red bars) or control missense-(white bars) infused animals (Sidak’s multiple comparisons test, *ns*).

#### Unilateral infusion of orexin antisense reduced orexin cell numbers

In rats with LH morpholino injection, we observed a significant reduction (~35%) in the number of orexin-expressing neurons in LH (Figure 2B,C; unpaired *t*-test; t_15_=3.55; *p*<0.01), but no change in orexin expression in Pef/DMH (Figure 2B,D; unpaired *t*-test; t_15_=0.49; *ns*). The number of orexin neurons in the uninjected hemisphere of morpholino-infused animals was similar to that observed in animals from Experiment 1, indicating no compensatory effects from morpholino treatment in the contralateral hemisphere (data not shown; unpaired *t*-test; t_19_=1.084; ns). Orexin-A morpholino antisense microinjections produced varying degrees of knockdown in these animals, depending on the location of cannula. We observed that antisense-infused animals with cannula targeted dorsal to the top of the fornix had a knockdown of ~25% of orexin cell numbers compared to the uninjected hemisphere, and this was statistically significant (n=5) (data not shown, unpaired *t*-test, t_8_= 2.69, p<0.05). This knockdown was proportional to the knockdown observed in our previous publication (James, Stopper, Zimmer et al., 2018) where injections were made bilaterally into the same dorsal region.

#### Unilateral infusion of antisense reduced motivation

A representative demand curve 6d following orexin-A morpholino antisense in LH unilaterally is compared to the baseline demand (prior to antisense injection; within-subject) in Figure 2E. Overall, unilateral orexin-A antisense into LH significantly increased α (decreased motivation) (Figure 2F, two-way RM ANOVA, main effect of treatment: F_1,16_= 16.61, *p*<0.001; interaction of treatment x morpholino type: F_1,16_=5.52, *p*<0.05). No changes were observed in Q_0_ for either orexin-A antisense or control missense-infused animals (Figure 2G, two-way RM ANOVA, *ns*). Of note, the change in α observed here following unilateral injections of morpholino was roughly half that reported in our previous study using bilateral LH orexin knockdown, and this trended towards significance (data not shown, paired samples t-test, t_4_=2.67, *p*=0.056).

#### The number of LH orexin neurons correlated with motivation for cocaine

Within antisense-infused animals, we observed a negative correlation between overall LH orexin cell numbers (summed across injected and non-injected hemispheres) and α values, such that animals with higher motivation (lower α values) had greater LH orexin+ neurons (Figure 3A, β =-0.62, *p*<0.05). In contrast, Q_0_ values were not significantly correlated with the number of LH orexin cells (Figure 3B, β= 0.41, *p*=0.09). No associations were observed between the number of Pef/DMH orexin neurons and α (Figure 3C, β=-0.35, *p*=0.41) or Q_0_ (Figure 3D, β=0.29, *p*=0.49).

**Figure 3.**
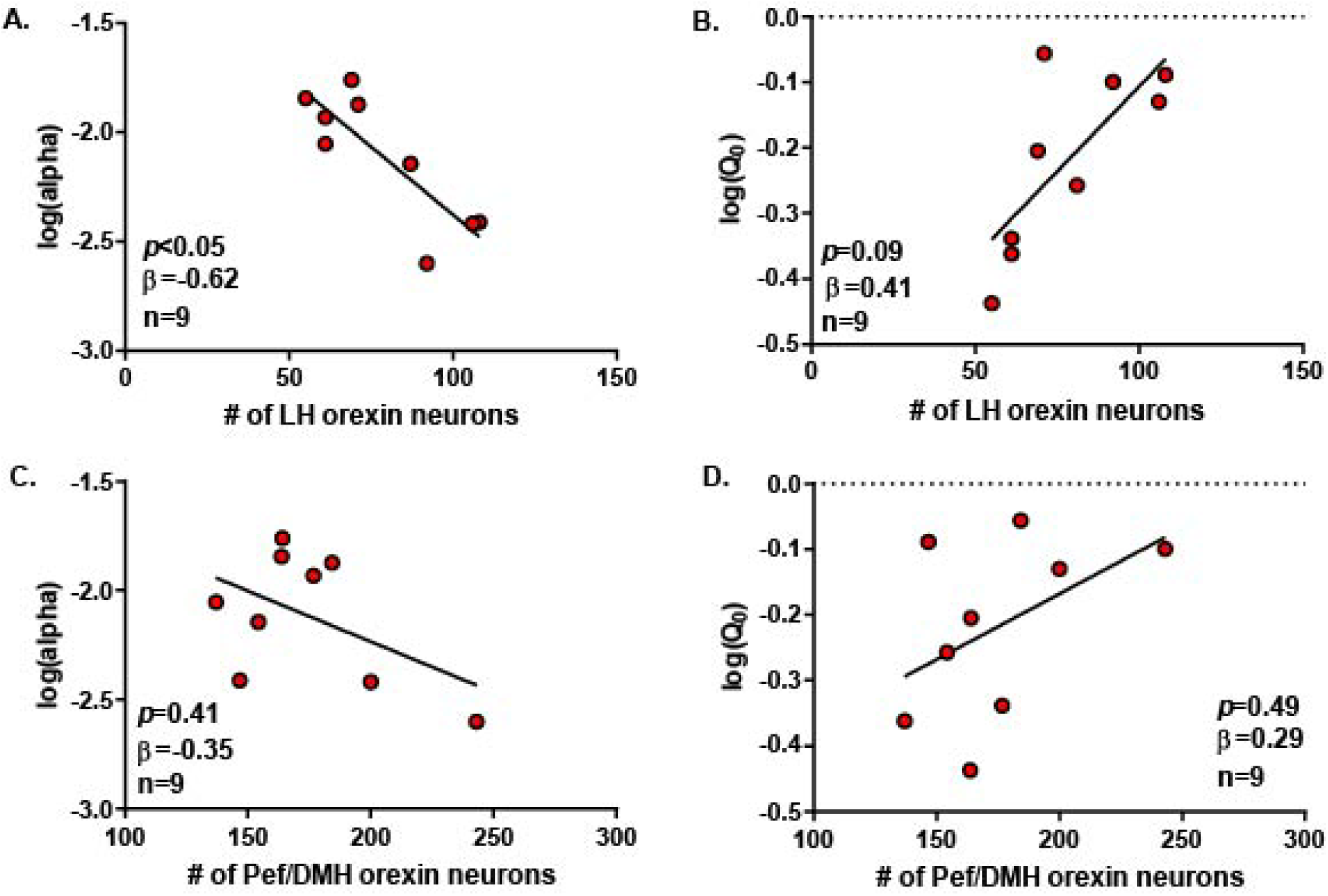
LH orexin neuron number predicts cocaine demand elasticity (α) and baseline consumption (Q_0_). <u>A.</u> The number of LH orexin neurons predicted α (multiple linear regression, *p*<0.05) but not <u>B.</u> Q_0_ values (multiple linear regression, *ns*). <u>C.</u> There was no relationship between Pef/DMH orexin cell number and α (multiple linear regression, *ns*) <u>D.</u> or Q_0_ values (multiple linear regression, *ns*).

We found that the percentage change in α correlated with the amount of LH orexin knockdown in antisense-injected animals when compared to the uninjected hemisphere, such that animals with a greater change in baseline α had a larger LH orexin cell knockdown (Figure 4A, β=-0.71, *p*<0.05). There was no such association between the change in Q_0_ and LH orexin knockdown (Figure 4B, β=0.09, *p*=0.78).

**Figure 4.**
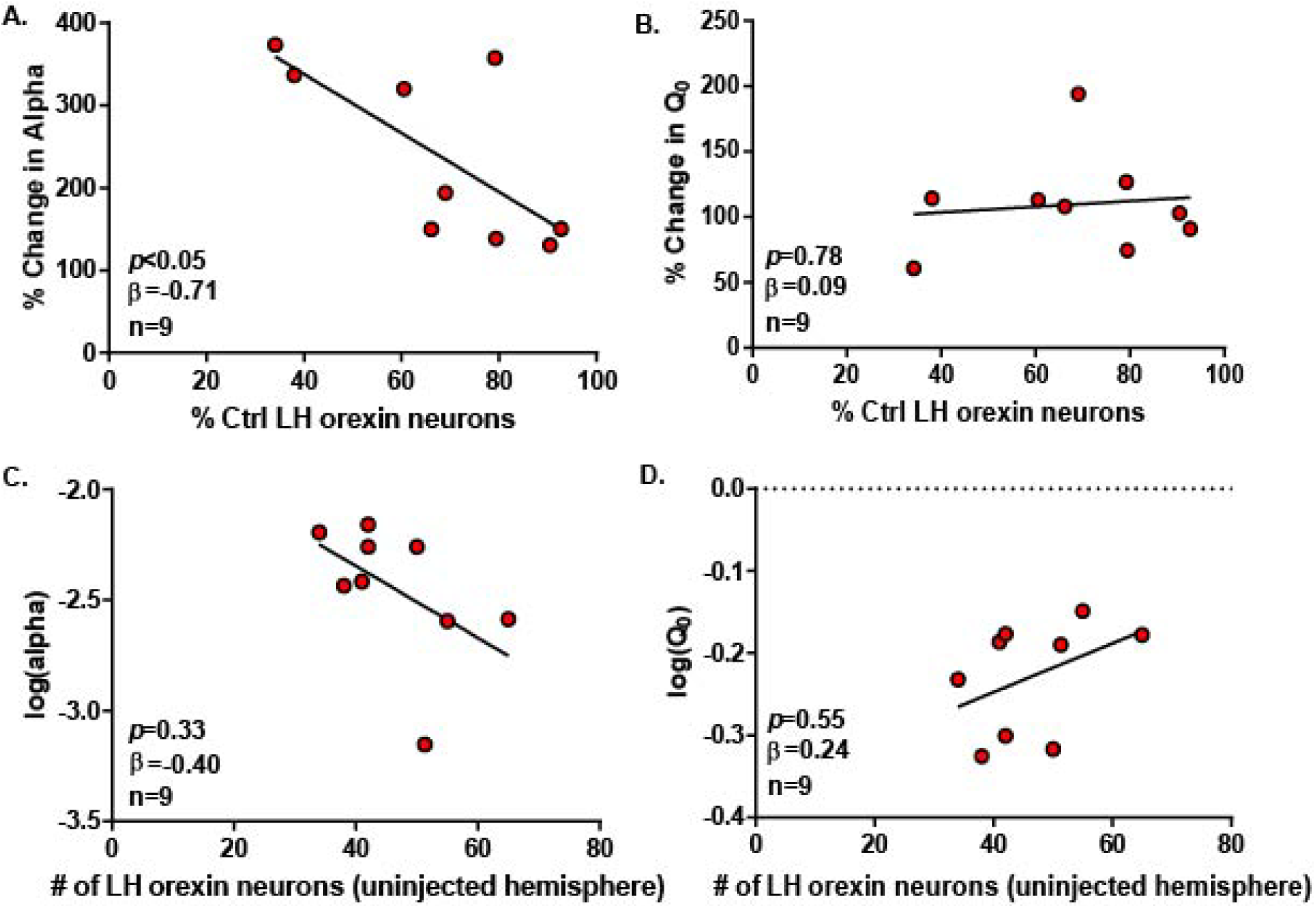
LH orexin neuron knockdown predicts antisense-induced changes in cocaine demand elasticity. <u>A.</u> Animals with greater LH orexin knockdown following antisense infusion, when orexin neurons in the injected hemisphere were compared to the uninjected hemisphere, had a larger change in α value (multiple linear regression, \**p*<0.05). <u>B.</u> The degree of LH orexin knockdown did not predict changes in Q_0_ post-antisense microinjection (multiple linear regression, *ns)*. <u>C.</u> In antisense-infused animals, there was a non-significant relationship between the numbers of LH orexin neurons in the uninjected hemisphere and baseline (pre-antisense injection) α values (multiple linear regression, *ns*). <u>D.</u> The number of LH orexin neurons did not predict baseline Q_0_ in the uninjected hemisphere (multiple linear regression, *ns*).

Finally, we correlated baseline (pre-morpholino injection) cocaine demand elasticity (α) with the number of orexin-expressing neurons in the uninjected hemisphere (as a measure of endogenous orexin levels). Consistent with Experiment 1, there was a strong trend towards a negative correlation between these two measures, although this failed to reach statistical significance (Figure 4C, β=-0.40, *p*=0.33). There was no association between baseline Q_0_ and LH orexin neurons in the uninjected hemisphere (Figure 4D, β=0.24, *p*=0.55).

### Experiment 3: Relationship between LH orexin cell number and demand following abstinence

To determine if the relationship between LH orexin expression and demand persisted after abstinence or with locomotor activity, we quantified LH and Pef/DMH orexin cells in BE-experienced animals sacrificed immediately after locomotor testing which followed 2 wk of abstinence from cocaine self-administration (n=8). LH orexin cell number inversely correlated with animals’ baseline α (measured 2 wk prior to sacrifice; Figure 5A, Pearson correlation, *p*<0.05), whereas Pef/DMH orexin expression did not (Figure 5B, Pearson correlation, *p*= 0.770). This demonstrates that LH orexin cell numbers during abstinence reflected prior individual differences in motivation for cocaine. We also observed that the number of LH orexin cells did not correlate with total locomotor activity in the test prior to sacrifice (Figure 5C, Pearson correlation, *p*=0.982). This reveals that individual differences in numbers of orexin-expressing neurons in Experiments 1 and 2 were not likely due to differences in general arousal/motor activity. We also found no difference in the number of LH orexin neurons in animals perfused following locomotor testing and animals Experiment 1 (Figure 5D, unpaired samples *t*-test, t_18_=0.123, *p*=0.903).

**Figure 5.**
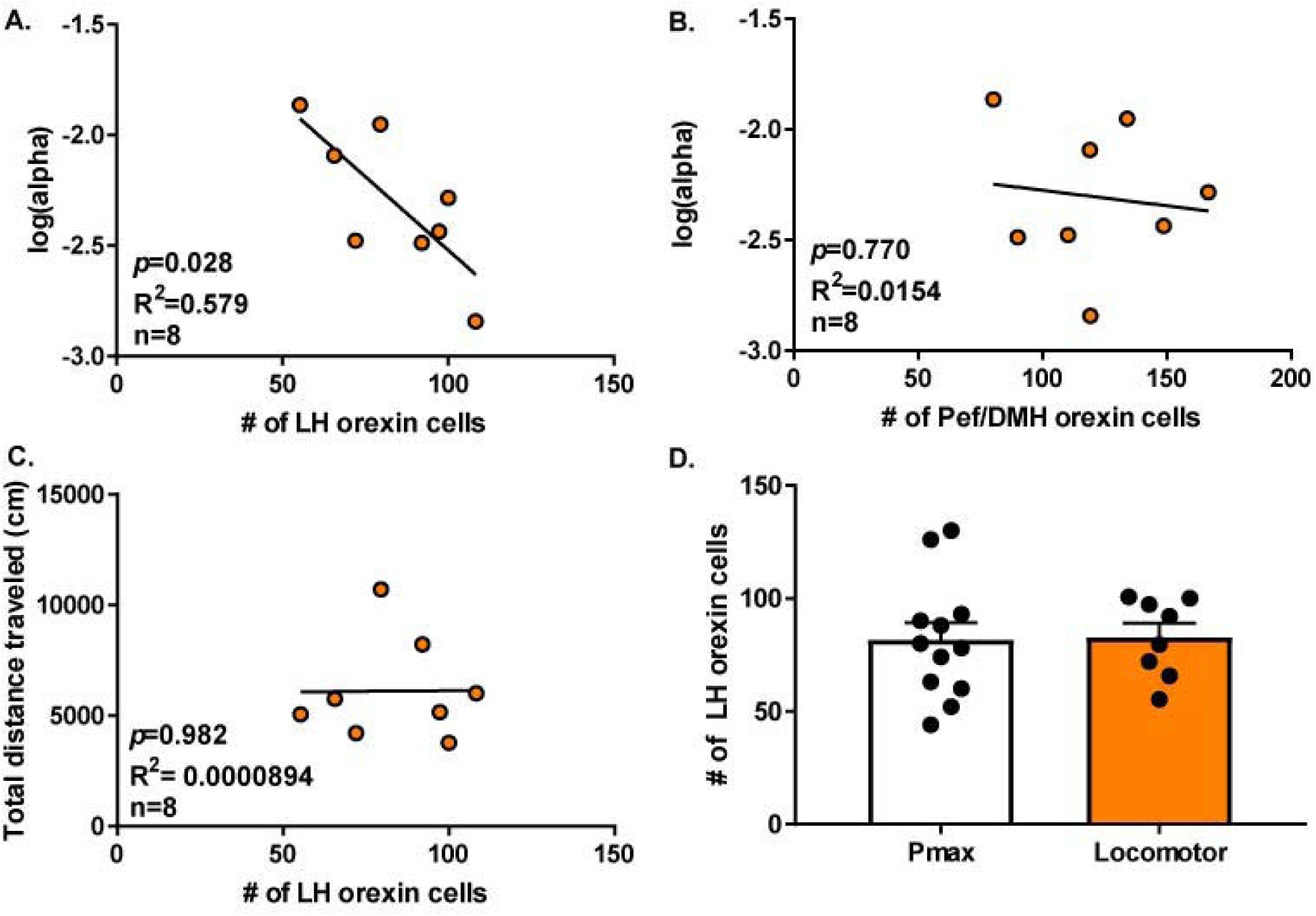
Numbers of LH orexin neurons predict individual differences in α values following abstinence from cocaine. A. LH orexin cell numbers at 2 weeks of cocaine abstinence correlated with animals’ baseline α values measured two weeks previously (Pearson correlation, *p*<0.05). B. No relationship was observed between baseline α values and the number of Pef/DMH orexin cells (Pearson correlation, *ns*). C. The number of LH orexin neurons in locomotor-tested animals did not predict habituated locomotor activity (Pearson correlation, *ns).* D. Animals sacrificed immediately following locomotor testing (locomotor) had similar numbers of LH orexin neurons compared to animals sacrificed 90 minutes post-Pmax (Pmax unpaired samples t-test, *ns*).

## Discussion

We show that the number of orexin cells in LH predicts motivation (demand elasticity) for cocaine. A similar relationship was observed when tissue was collected from rats that underwent 2wk abstinence from cocaine. Differences in LH orexin number were not due to variability in arousal state, as orexin expression did not correlate with general locomotor activity. In addition, knockdown of orexin signaling reduced demand (α) but not low effort consumption of cocaine (Q_0_), and degree of knockdown correlated with the change in α following knockdown, consistent with other studies from our lab (Bentzley & Aston-Jones, 2015; James, Stopper, Zimmer et al., 2018). These studies are consistent with the view that orexin mediates the motivational properties of cocaine without influencing the hedonic value of the drug (James, Mahler, Moorman et al., 2017; Mahler, Moorman, Smith et al., 2014). Taken together, our results indicate that the number of orexin-expressing neurons in LH is a determinant of individual differences in cocaine motivation.

### High motivation animals have greater orexin expression in lateral hypothalamus

Across all three experiments, animals with more LH orexin cells had higher motivation for cocaine. In Experiment 1, high motivation (low demand elasticity) animals had greater LH orexin expression than low motivation animals (high demand elasticity) animals. In Experiment 2, the number of remaining LH orexin cells following orexin-A morpholino antisense knockdown correlated with demand elasticity for cocaine, and the change in LH orexin cell number correlated with the change in α value. The relationship between demand elasticity and LH orexin neuron number does not require the presence of the drug or drug-paired context because in Experiment 3, the number of LH orexin cells in animals sacrificed immediately after general locomotor testing correlated inversely with cocaine demand elasticity 2 wk prior. Although other studies reported that chronic exposure to drugs of abuse upregulated numbers of LH orexin-expressing cells (James, Stopper, Zimmer et al., 2018; Lawrence, Cowen, Yang et al., 2006; Thannickal, John, Shan et al., 2018), to our knowledge our results are the first to show a relationship between the endogenous number of LH orexin cells and baseline addiction propensity. Taken together, these results are consistent with our hypothesis that orexin signaling is particularly important in highly motivated individuals (James, Bowrey, Stopper et al., 2018), and that the number of LH orexin neurons could serve as a biomarker of addiction susceptibility.

We proposed that orexin signaling translates motivational drive into behavioral output (Mahler, Moorman, Smith et al., 2014), particularly in response to drug-associated cues and contexts (Bentzley & Aston-Jones, 2015; Smith, See & Aston-Jones, 2009; Smith, Tahsili-Fahadan & Aston-Jones, 2010; Zhou, Ghee, Chan et al., 2012). Our lab has shown that LH orexin neurons in particular mediate reinstatement of drug seeking elicited by drug-associated stimuli (Harris, Wimmer & Aston-Jones, 2005; James, Mahler, Moorman et al., 2017; Mahler, Smith, Moorman et al., 2012; Sartor & Aston-Jones, 2012). Thus, higher expression and/or activation of these cells may attach greater motivational significance to cocaine-associated stimuli and result in greater motivation (demand) for cocaine. A recent study in hypocretin/orexin knockout mice demonstrated that animals lacking the neuropeptide have reduced incubation of cocaine craving and do not reinstate drug seeking in response to cocaine-paired cues (Steiner, Rossetti, Sakurai et al., 2018). Greater numbers of cells promoting cue reactivity in high motivation animals may also explain our previous finding that animals with low α measured in BE have greater cue-induced reinstatement of cocaine seeking (Bentzley, Jhou & Aston-Jones, 2014). Therefore, a greater number of LH orexin-expressing cells may result in higher cue reactivity and greater drug seeking in highly motivated animals.

Although we did not directly assess the pathways through which orexin mediates motivation for cocaine, other studies indicate that VTA is likely an important target region. Significant data indicates that orexin neurons drive motivational processes via projections to ventral tegmental area (VTA) (Mahler, Moorman, Smith et al., 2014). Orexin potentiates cocaine-induced plasticity of VTA DA neurons *in vitro* (Borgland, Storm & Bonci, 2008; Borgland, Taha, Sarti et al., 2006), and orexin-glutamate interactions in VTA are necessary for cue-associated cocaine seeking (Mahler, Smith & Aston-Jones, 2013). Blocking orexin-1 receptor signaling in VTA reduces cocaine seeking (James, Charnley, Levi et al., 2011; Mahler, Smith & Aston-Jones, 2013) or effortful self-administration (Espana, Oleson, Locke et al., 2010), whereas infusion of orexin A into VTA reinstates extinguished cocaine seeking (Harris, Wimmer & Aston-Jones, 2005; Wang, You & Wise, 2009). Orexin input to VTA increases DA release in nucleus accumbens (NAc) (Baimel, Lau, Qiao et al., 2017; Espana, Melchior, Roberts et al., 2011; Prince, Rau, Yorgason et al., 2015), which is associated with goal-directed behavior (Saddoris, Sugam, Cacciapaglia et al., 2013). Greater LH orexin input to the mesolimbic dopamine system may drive greater cocaine taking in high motivation animals, and future experiments could seek to determine if greater LH orexin input to VTA circuit specifically mediates motivation for cocaine.

Consistent with a selective role for the orexin system in motivated responding for cocaine, we generally saw no relationship between orexin cell numbers and low-effort cocaine intake. We did not observe a difference in the number of orexin cells between high and low takers (high and low Q_0_) in Experiment 1. Also, we observed no change in Q_0_ following antisense injections (Experiment 2), indicating no causal relationship between orexin cell numbers and low-effort cocaine intake. These findings are consistent with previous studies from our lab and others indicating that blockade of orexin-1 receptor signaling has no effect on low effort (FR1) cocaine consumption (Borgland, Chang, Bowers et al., 2009; Espana, Melchior, Roberts et al., 2011; Espana, Oleson, Locke et al., 2010; Hollander, Pham, Fowler et al., 2012; Smith, See & Aston-Jones, 2009).

### Unilateral LH orexin knockdown is sufficient to reduce cocaine demand

Unilateral orexin knockdown with orexin-A morpholino antisense was sufficient to reduce motivation for cocaine (increase demand elasticity), but not low effort consumption (Q_0_). Peak selective knockdown of orexin A protein occurs six days post-antisense without affecting expression of interdigitated neurons containing melanin-concentrating hormone (James, Stopper, Zimmer et al., 2018; Reissner, Sartor, Vazey et al., 2012; Sartor & Aston-Jones, 2012). Although other studies have shown no effect of unilateral orexin inhibition or knockdown on behavior (Mahler, Smith & Aston-Jones, 2013; Sartor & Aston-Jones, 2012), ours is the first to examine unilateral orexin knockdown during BE performance requiring high levels of motivation. It may be that orexin neurons bilaterally project to reward-associated areas like VTA, and unilateral knockdown reduces orexin input to both hemispheres the target brain area. Orexin collateralization in reward-associated areas has largely been unexplored. Thus, future anatomical studies should investigate the extent to which orexin inputs collateralize in reward-associated regions like VTA.

### Lateral hypothalamus orexin neuron number correlates with motivational and not arousal state

Our findings revealed that the number of orexin neurons in LH, but not in Pef/DMH, predicted motivation for cocaine as assessed by the BE paradigm (demand elasticity). Our lab also recently determined that knockdown of orexin in LH, but not in Pef/DMH, reduced motivation for cocaine (James, Stopper, Zimmer et al., 2018). We also observed that the number of LH orexin neurons did not correlate with locomotor activity, indicating that differences in cell number were not related to non-specific changes in, e.g., arousal state. Taken together, these results support our lab’s hypothesis that LH orexin cells are preferentially involved in regulating motivational processes, and are less important for arousal/stress regulation (Aston-Jones, Smith, Sartor et al., 2010; Harris & Aston-Jones, 2006; James, Mahler, Moorman et al., 2017).

Our results are consistent with several studies from our lab and others indicating that individual differences in the LH orexin system are associated with reward/motivation levels. LH orexin neurons and mRNA expression is upregulated in animals with high motivation for drug (James, Stopper, Zimmer et al., 2018; Lawrence, Cowen, Yang et al., 2006; Thannickal, John, Shan et al., 2018). In addition, Fos expression specifically in LH orexin cells correlates with preference for both morphine and cocaine (Harris, Wimmer & Aston-Jones, 2005; Harris, Wimmer, Randall-Thompson et al., 2007; Lasheras, Laorden, Milanes et al., 2015; Richardson & Aston-Jones, 2012; Sartor & Aston-Jones, 2012). The preferential involvement of LH orexin cells in drug seeking may be mediated by distinct inputs or outputs compared to Pef/DMH cells (Fadel & Deutch, 2002; Gonzalez, Jensen, Fugger et al., 2012), though anatomical evidence thus far remains unclear.

Collectively, these studies demonstrate that the number of LH orexin neurons is a significant factor contributing to individual differences in motivation for cocaine and may be a previously unrecognized but important element in addiction pathology.

